# Response to Therapeutic Sleep Deprivation: A Naturalistic Study of Clinical and Genetic Factors and Post-Treatment Depressive Symptom Trajectory

**DOI:** 10.1101/179457

**Authors:** Nina Trautmann, Jerome C. Foo, Josef Frank, Stephanie H. Witt, Fabian Streit, Jens Treutlein, Steffen Conrad von Heydendorff, Maria Gilles, Major Depressive Disorder Working Group of the Psychiatric Genomics Consortium, Andreas J. Forstner, Ulrich Ebner-Priemer, Markus M. Nöthen, Michael Deuschle, Marcella Rietschel

## Abstract

Research has shown that therapeutic sleep deprivation (SD) has rapid antidepressant effects in the majority of depressed patients. Investigation of factors preceding and accompanying these effects may facilitate the identification of the underlying biological mechanisms. This exploratory study aimed to examine clinical and genetic factors predicting response to SD and determine the impact of SD on illness course. Mood and tiredness during SD were also assessed via visual analogue scales (VAS). Depressed inpatients (n = 78) and healthy controls (n = 15) underwent ~36hrs of SD. Response to SD was defined as a score of ≤2 on the Clinical Global Impression Scale for Global Improvement. Depressive symptom trajectories were evaluated for up to a month using self/expert ratings. Impact of genetic burden was calculated using polygenic risk scores for major depressive disorder. 72% of patients responded to SD. Responders and nonresponders did not differ in baseline self/expert depression symptom ratings, but mood subjectively measured by VAS scale differed. Response was associated with lower age (*p* = 0.007) and later age at life-time disease onset (*p* = 0.003). Higher genetic burden of depression was observed in non-responders than healthy controls. Up to a month post-SD, depressive symptoms decreased in both patients groups, but more in responders, in whom effects were sustained. The present findings suggest that re-examining SD with a greater focus on biological mechanisms will lead to better understanding of mechanisms of depression.

## Introduction

Therapeutic sleep deprivation (SD) reliably induces rapid and substantial antidepressant effects in the majority of patients with a major depressive episode (Benedetti *et al*, 2007; Benedetti and Colombo, 2011; Gillin, 1983; Wu and Bunney, 1990). A recent meta-analysis of SD studies showed an average response rate of approximately 50% with significant variability, with up to 78% of patients responding to SD treatment (Boland *et al*, 2017). Although its therapeutic value is limited due to relapse after recovery sleep (Giedke and Schwärzler, 2002; Wu *et al*, 1990), SD is particularly unique in its defined immediate positive effect on depressive mood and may therefore offer unique insights about the biological factors underlying depression. Response to SD has been associated with various factors, including circadian rhythms (Haug, 1992; Martiny *et al*, 2013; Reinink *et al*, 1990; Szuba *et al*, 1991); tiredness (Bouhuys *et al*, 1995); disease diagnosis and “endogenous depression” (Barbini *et al*, 1998; Benedetti *et al*, 2005; Larsen *et al*, 1976; Pflug and Tölle, 1971b); age-related features (Clark and Golshan, 2007; Cole and Muller, 1976; Pflug and Tölle, 1971a; Rudolf and Tölle, 1978) and candidate gene variants (Benedetti *et al*, 2011; Benedetti *et al*, 1999; Bunney *et al*, 2015). Although several plausible hypotheses have been formulated (Borbély *et al*, 2016; Borbely and Wirz-Justice, 1982; Bunney and Bunney, 2013; Wolf *et al*, 2016), a comprehensive understanding of underlying factors, especially with respect to the biological mechanisms involved, has not yet been achieved.

MDD is a heterogeneous disorder, and it is thought that a multitude of genetic variants are involved in course, development and response to treatment (Flint and Kendler, 2014; Sullivan *et al*, 2000). Understanding the role of genetic risk in modulation of response to treatment might allow the identification of potential responders, eventually leading to improvements in personalized care. It has been observed that higher genetic burden for psychiatric disorders is associated with response to treatment (Amare *et al*, In press; Frank *et al*, 2015; Tansey *et al*, 2013).

Recent genome-wide association studies with large samples have made substantial progress with identification of common risk variants for MDD (Wray and Sullivan, 2017). Furthermore, polygenic risk scores (PRSs), which summarize the effects of many single nucleotide polymorphisms (SNPs) in a single risk score offer the ability to associate burden of disease with clinical and phenotypic factors, and have been successfully applied to explore the genetic architecture of complex disorders (Frank *et al*, 2015; Schizophrenia Working Group of the Psychiatric Genomics, 2014; Wray *et al*, 2014).

In this naturalistic exploratory study, we assessed clinical and genetic factors associated with response to SD, going beyond the study of individual candidate genes for the first time, using all-genomic information in the form of PRSs. We also evaluated mood and tiredness longitudinally during SD, and the impact of SD on the further trajectory of depressive symptoms.

## Materials and Methods

### Participants

Seventy-eight inpatients (34 females; age mean ± standard deviation = 43.54 ± 14.80 years) presenting with an episode of major depression (unipolar, n = 71; bipolar I, n = 6; and bipolar II, n = 1) participated in this study. Depression was diagnosed according to ICD-10 criteria. Patients were recruited between August 2013 and April 2015 from consecutive admissions to the depression unit of the Central Institute of Mental Health (CIMH) in Mannheim, Germany. The study protocol stipulated that for 5+ days prior to SD, no changes were allowed to the medication regimen. Prescribed medication included typical and atypical antidepressants, lithium, and adjunct therapies (for details, see supplementary text). Fifteen healthy controls (8 females; 40.53 ± 15.90 years) with no history of psychiatric/somatic disorders were recruited through an online advertisement on the CIMH website. The investigation was carried out in accordance with the latest version of the Declaration of Helsinki and approved by the local ethics committee. All participants provided written informed consent following a detailed explanation of the study.

### Sleep Deprivation

Participation began on Day 1 (see Figure 1 schematic for details) whereupon baseline variables (see below) were assessed. During Day 2, patients engaged in normal ward routines.

**Fig.1.**
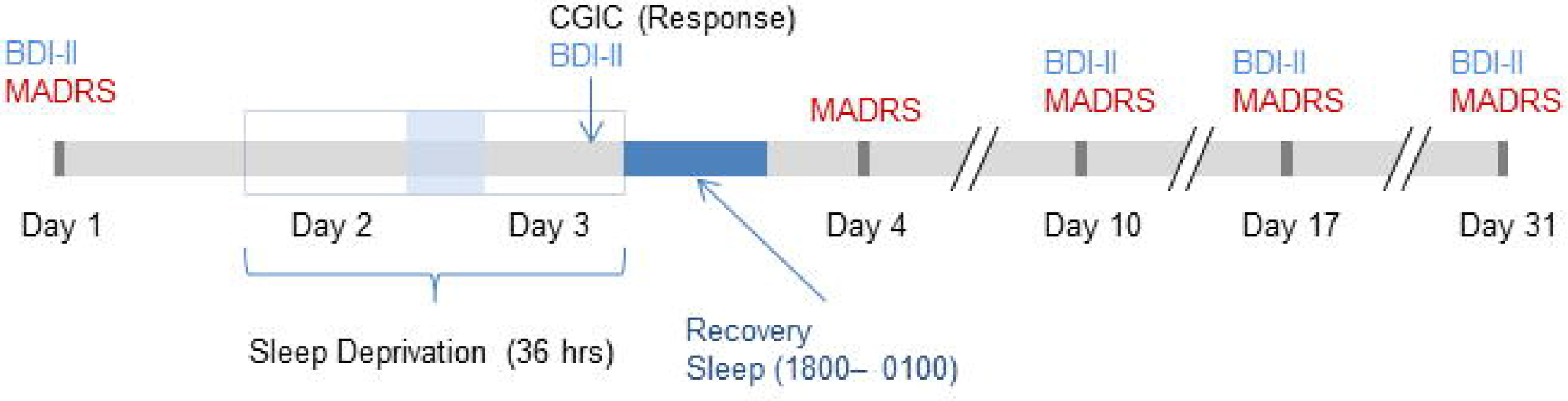
Schematic timeline of study schedule. CGI = Clinical Global Impression; BDI-II = Beck Depression Inventory II; MADRS = Montgomery- Äsberg Depression Rating Scale.

**Fig.2.**
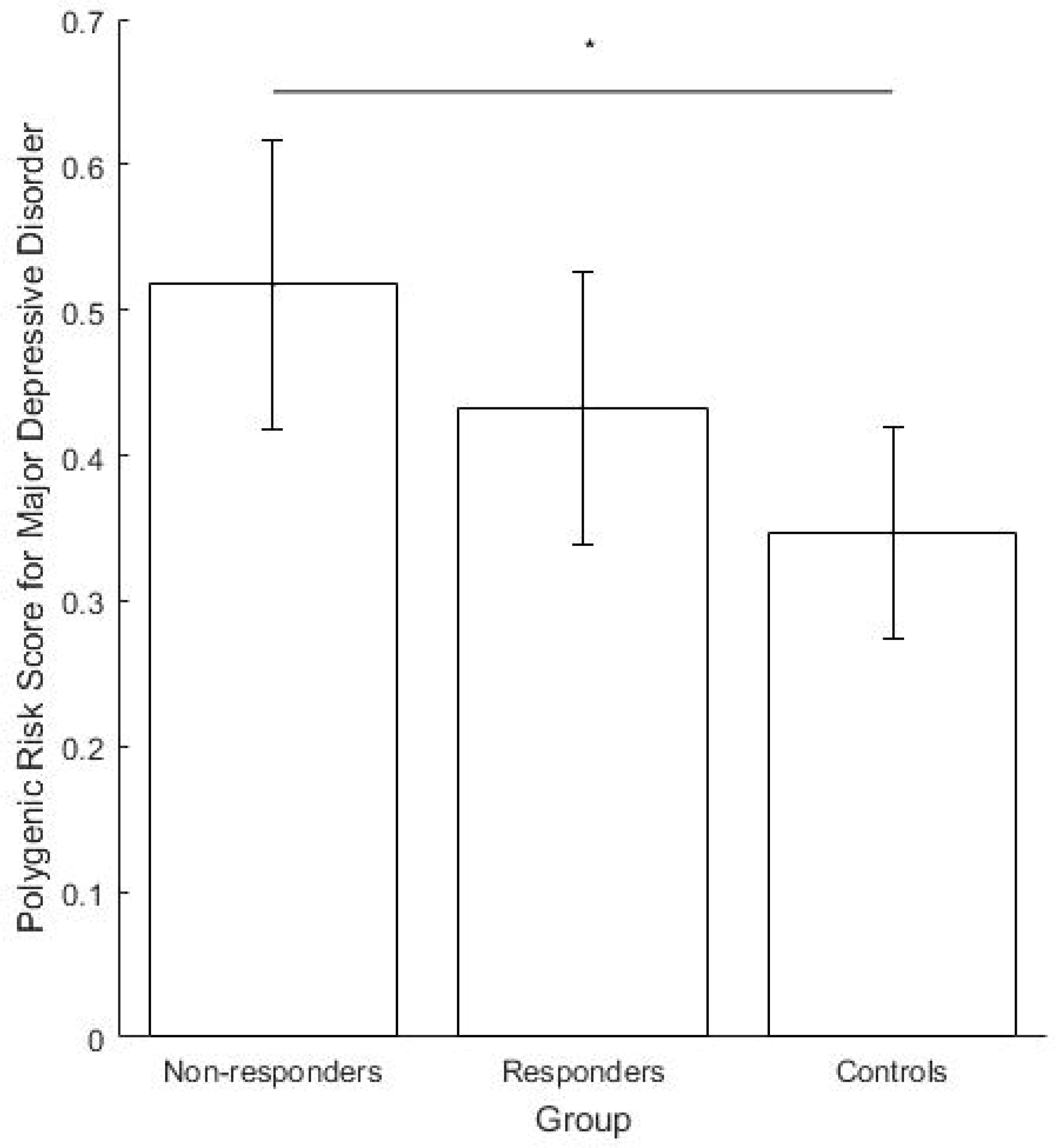
Polygenic Risk Scores (PRS) for major depression in non-responders, responders and healthy controls. Scores are normalised to 0-1. Error bars denote 95% confidence intervals. * *p* < 0.05.

SD was conducted in small groups of 1-5 participants under staff supervision. Participants remained awake from ~0600hrs on Day 2 to 1800hrs on Day 3 (36 hours). On Day 3, patients engaged in normal ward routines until undergoing recovery sleep from 1800-0100hrs. Sleep phase advance was then carried out, shifting sleep one hour forward each day until the patient’s regular sleep pattern was reached. Controls underwent SD alongside patients; their participation ended after the first recovery sleep.

### Data collection

#### Blood sampling

On Day 1, a venous blood sample was collected from participants for genome-wide genotyping, which was performed using the Global Screening Array (Illumina, Inc., San Diego, CA, USA). Genotyping and quality control procedures are described in detail in the supplement and elsewhere (Frank *et al*, 2015; Schizophrenia Working Group of the Psychiatric Genomics, 2014).

#### Demographic and clinical characteristics

On Day 1, the following factors were assessed: demographics, including *sex, age, age at initial disease onset (AaO);* clinical parameters (*body mass index [BMI], pulse);* history of psychiatric and somatic disorders and *family history* (*FH*) of MDD or bipolar disorder (BD).

The validated German version (Griefahn *et al*, 2001) of the Morningness-Eveningness-Questionnaire (D-MEQ) (Horne and Ostberg, 1976) was used to assess circadian rhythm/diurnal variation. The D-MEQ comprises 19 items on circadian patterns, identifying *morning, intermediate*, and *evening* chronotypes.

#### Response to SD

*Response* to SD was evaluated between 1600-1700hrs on Day 3 by the senior clinical researcher (MD) using the Clinical Global Impression Scale for Global Improvement (CGIC) (Guy, 1976). Possible CGIC scores were: 1 = Very much improved; 2 = Much improved; 3 = Minimally improved; 4 = No change; 5 = Minimally worse; 6 = Much worse; 7 = Very much worse. Response and non-response to SD were defined as scores of ≤ 2 and ≥ 3, respectively.

#### Depressive Symptoms Scales

The 10-item Montgomery-Åsberg Depression Rating Scale (MADRS) (Montgomery and Asberg, 1979) was completed by the senior clinical researcher (MD) on Days 1, 4, 10, 17, and 30. The 21-item Beck Depression Inventory-II (BDI-II) (Beck *et al*, 1996) was completed by patients on Days 1, 3, 10, 17, and 30.

#### During SD: Visual Analogue Scale for Mood and Tiredness

Visual analogue scales (VAS) (Aitken, 1969) for *mood* and *tiredness* were completed by participants every two hours from 1000hrs on Day 2–1800hrs on Day 3. For both scales, values ranged from 0-10. *Mood* ratings ranged from: “worst mood imaginable” to “best mood imaginable”. *Tiredness* ratings ranged from: “not tired at all” to “so tired that it’s hard to stay awake”. Locomotor activity was acquired using the SOMNOwatch (SOMNOmedics GmbH, Germany), and patients recorded in a wear log when the device was worn/removed; these were inspected to identify subjects who had fallen asleep before assessment of response.

### Data Analysis

Statistical analyses were performed using IBM SPSS Statistics for Windows version 24. Statistical significance was set at *p* < 0.05.

#### Descriptive statistics

Descriptive statistics were calculated for all participants. For continuous variables, mean values were compared using independent samples t-tests. For nominal values, proportions were compared using Fisher’s exact test.

#### Genotyping and polygenic risk score calculation

Polygenic risk scores (PRS) (Wray *et al*, 2014) were calculated using genome-wide association data from the Psychiatric Genomics Consortium MDDII (Cases: n = 59851, Controls: n = 113154) (Wray *et al*, 2017). A *p*-value threshold of 1.0 was found to give best-fit (for details, see supplementary text). Scores were normalised to 0-1. Binomial logistic regression was used to compare PRS across disease state. To compare PRS across groups (non-responder/responder/control) one-way analysis-of-variance (ANOVA) was used.

#### Baseline predictors of response to SD

To identify baseline predictors of *response to* SD, a binomial logistic regression analysis was performed. *Response* was specified as the dependent variable. Categorical independent variables comprised: *sex; diurnal variation* (*morning/intermediate/evening chronotype*); *season (spring/summer/autumn/winter*); *diagnosis (Unipolar MDD/BD);* and *FH*. Continuous independent variables comprised: *PRS for MDD*; *age; AaO;* and baseline *BDI-II* and *MADRS* scores.

#### Mood and Tiredness Trajectories

To compare mood and tiredness trajectories between responders and non-responders during SD, a random-intercepts mixed model was used (accounting for intra-individual clustering of observations). *Mood and Tiredness* were specified as dependent variables in separate models. *PRS for MDD, Response* and *Timepoint* and the interaction between *Response*\**Timepoint* were specified as fixed factors. *Timepoint* was centred to midnight and included in a repeated term with an AR1 covariance structure.

#### Depressive Symptoms Score Trajectories

Correlations between *MADRS* and *BDI-II* scores were examined over all measurement days. Both score trajectories were examined using random-intercepts mixed models. Fixed effects included *sex, season, diagnosis, response*, and *measurement day* entered as factors. Age, *AaO and PRS for MDD* were entered as covariates. The *response*\**measurement day* interaction was entered as a fixed effect. *Measurement day* was included in a repeated term with a diagonal covariance structure.

## Results

### Demographics and descriptive statistics

Descriptive statistics are shown in **Table S1**. Six patients were excluded from the analysis as they did not complete SD. Four patients were excluded for having fallen asleep prior to response rating. Thus data from a total of 68 patients were included in the subsequent analyses (except for PRS analysis). A total of 49 (49/68; 72.1%) responded to SD.

### Polygenic Risk Scores

The regression model comparing PRS for disease state (controls n = 15; patients n = 72) found higher PRS in patients at the trend level (*p* = 0.068, ΔNagelkerke R^2^ = 0.066). The ANOVA to compare groups (responders n = 46, non-responders n = 18, controls n = 15) found a significant difference between groups (*F*_2,76_ = 3.426, *p* = 0.038). A post-hoc Tukey test found the group difference to be driven by higher scores in non-responders than controls (significant, *p* = 0.029). Although not significant, higher scores were found in non-responders than responders (*p* = 0.212) and controls than responders (*p* = 0.309) (see Figure 1 and supplementary text for additional details).

### Baseline Predictors of Response to SD

The regression model included 57 patients due to missing (assessment or genetic) data (**Table S2**). The model was statistically significant, *χ*^2^(13) = 24.477, *p* = 0.027, explaining 50.2% of the variance in response. Lower *age* (*p* = 0.007) and higher *AaO* (*p* = 0.003) were significantly associated with an increased likelihood of *response*. No significant effects were found for *PRS* (*p* = 0.907); *FH* (*p* = 0.125); *sex* (*p* = 0.148); *season (*p* =* 0.587); baseline *BDI-II score* (*p* = 0.986); baseline *MADRS score* (*p* = 0.314); *diagnosis* (*p* = 0.691); or *diurnal variation* (*p* = 0.343).

### Mood and Tiredness

Figure 3 shows the trajectories during SD for group mean *mood* (3a) and *tiredness* (3b) throughout SD.

**Fig.3.**
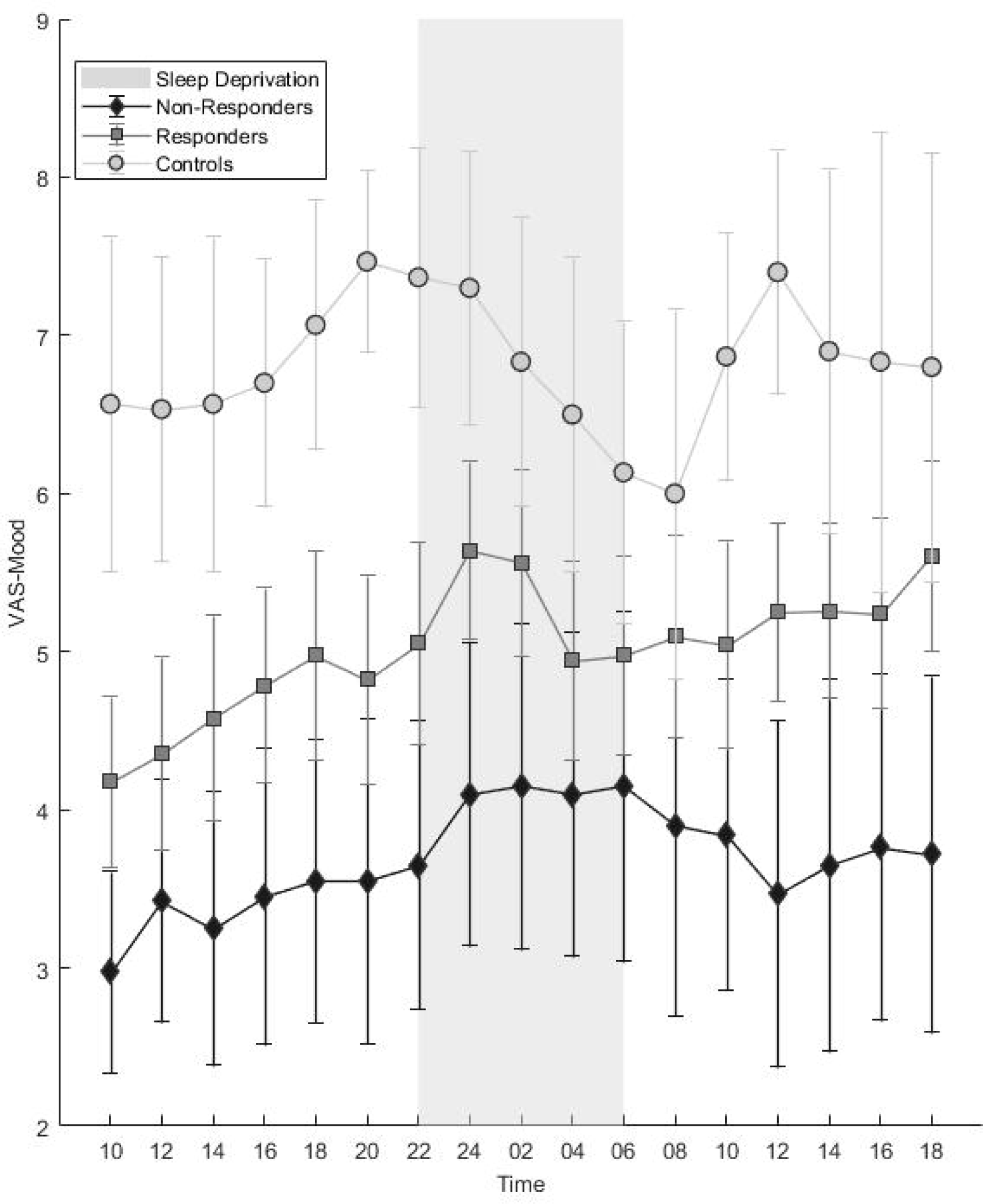
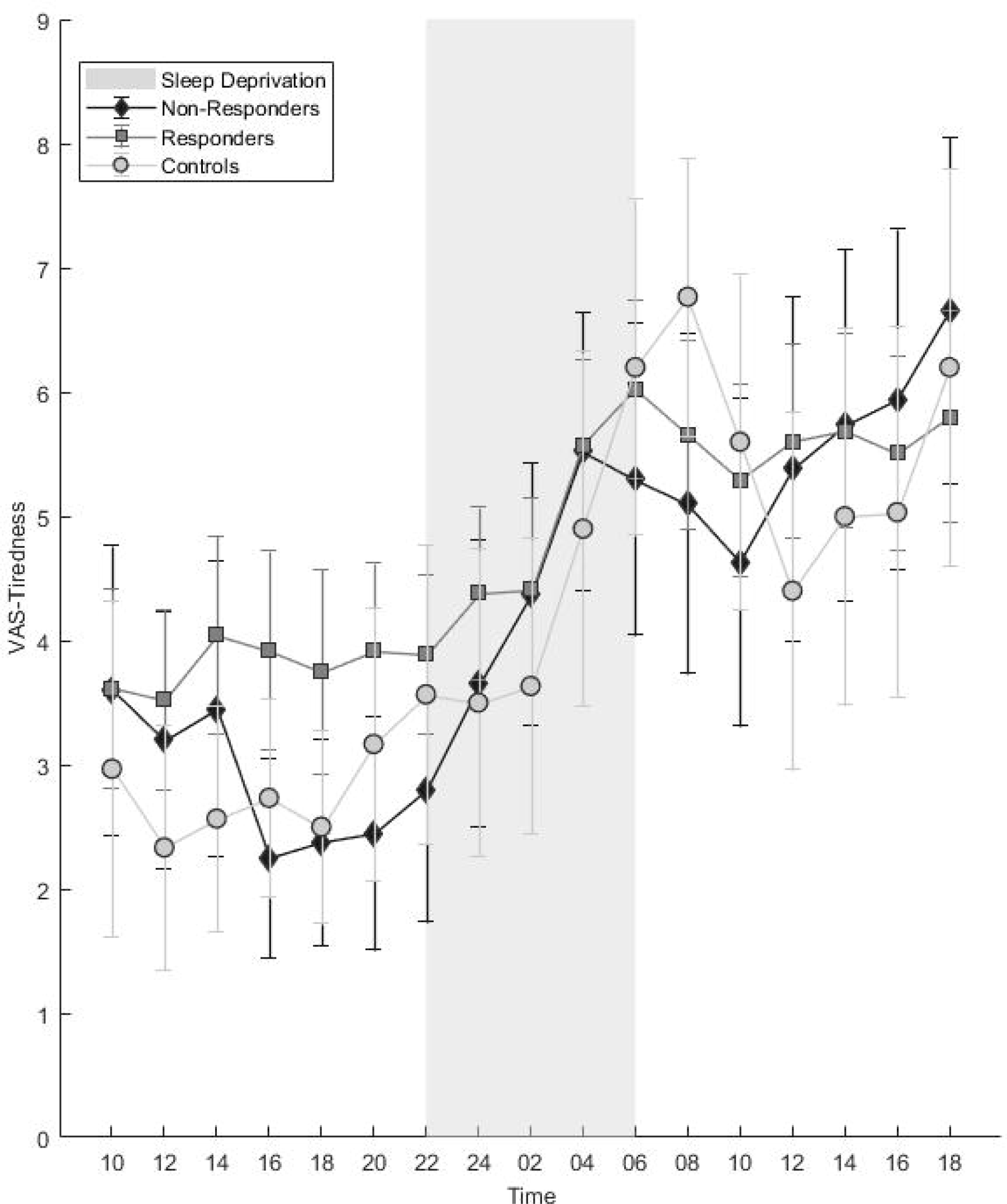
Trajectories of mean a) mood and b) tiredness during sleep deprivation. Error bars denote 95% confidence intervals.

In the linear mixed model analysis of mood (**Table S3a**), significant main effects of *timepoint* (*F*_16,540.801_ = 2.518, *p* = 0.001) and *response* (*F*_1,63.217_ = 8.811, *p* = 0.004) were observed. In the whole cohort, *mood* improved over *time* (**see Table S3a**) while worse mood was observed in nonresponders vs. responders (*t* = −2.109, *df* = 215.848, *p* = 0.036). No significant effects of interaction between *response*\**timepoint* were observed (*p* = 0.781; only at the final observation point did the interaction show a trend towards significance, *p* = 0.098). No significant association was observed with PRS (*p* = 0.276).

In the analysis of tiredness (**Table S3b**), a significant effect of *timepoint* (F_16,544.059_ = 11.662, *p* < 0.001) was observed; participants became increasingly tired as time progressed. No significant effect of *response* (*p* = 0.542) or overall *response*\**timepoint* interaction (*p* = 0.355) was observed, but examining the interaction term revealed lower tiredness scores in non-responders than responders at times 1600 (*p* = 0.040), 1800 (*p* = 0.061) and 2000 (*p* = 0.048). No significant association was observed with PRS (*p* = 0.389).

For both models, estimated correlation between any two consecutive assessment points was significant (AR1 rho, *p* < 0.001)

### Depressive Symptoms (MADRS and BDI-II)

Responders and non-responders did not differ in terms of baseline *MADRS* and *BDI-II* scores (Figure 4a,b). **Table S4a** shows the correlation between *MADRS* and *BDI-II* scores on all measurement days which was consistent (all Pearson r ≥ 0.4) and significant (all *p* < 0.001).

**Fig 4.**
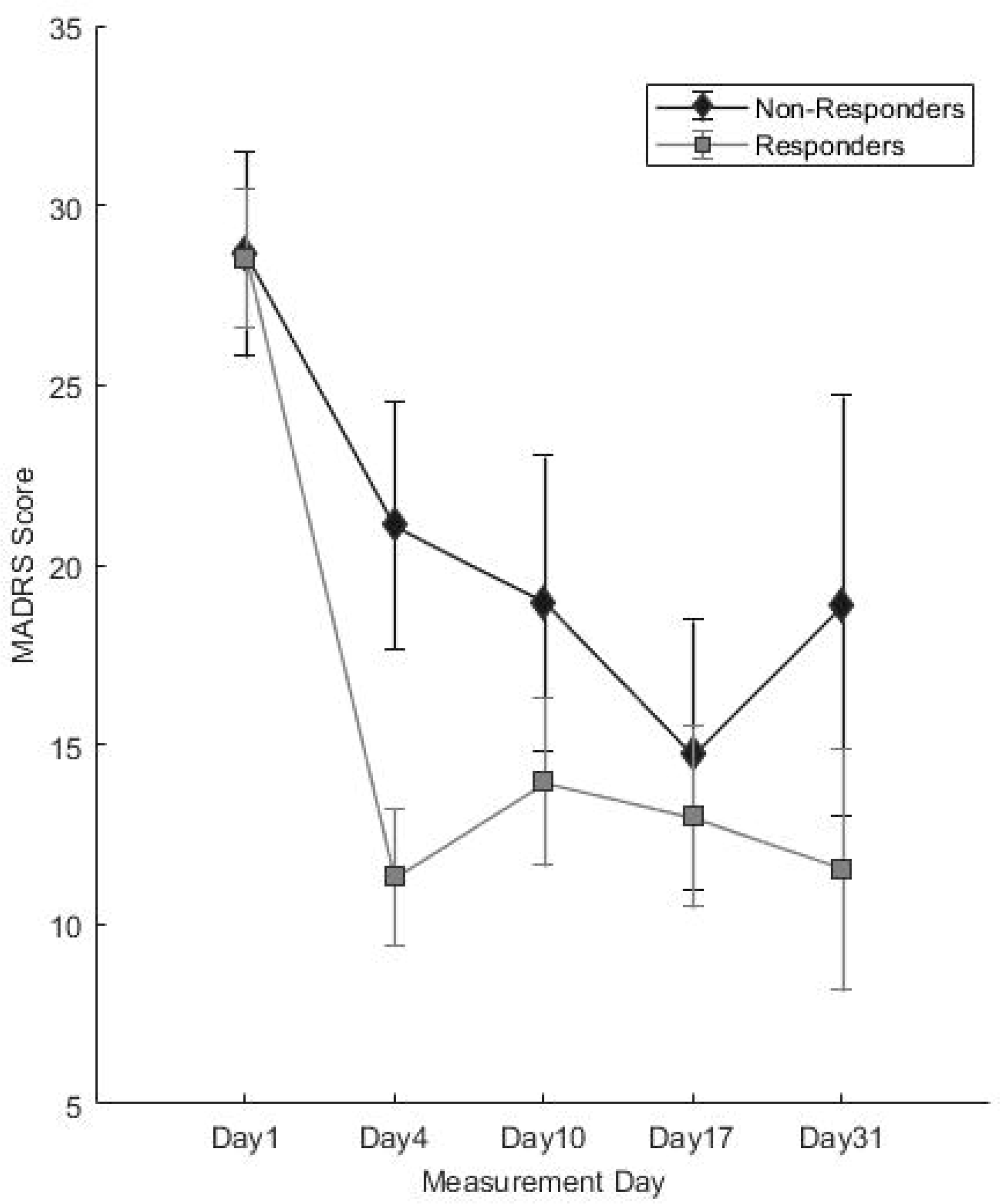
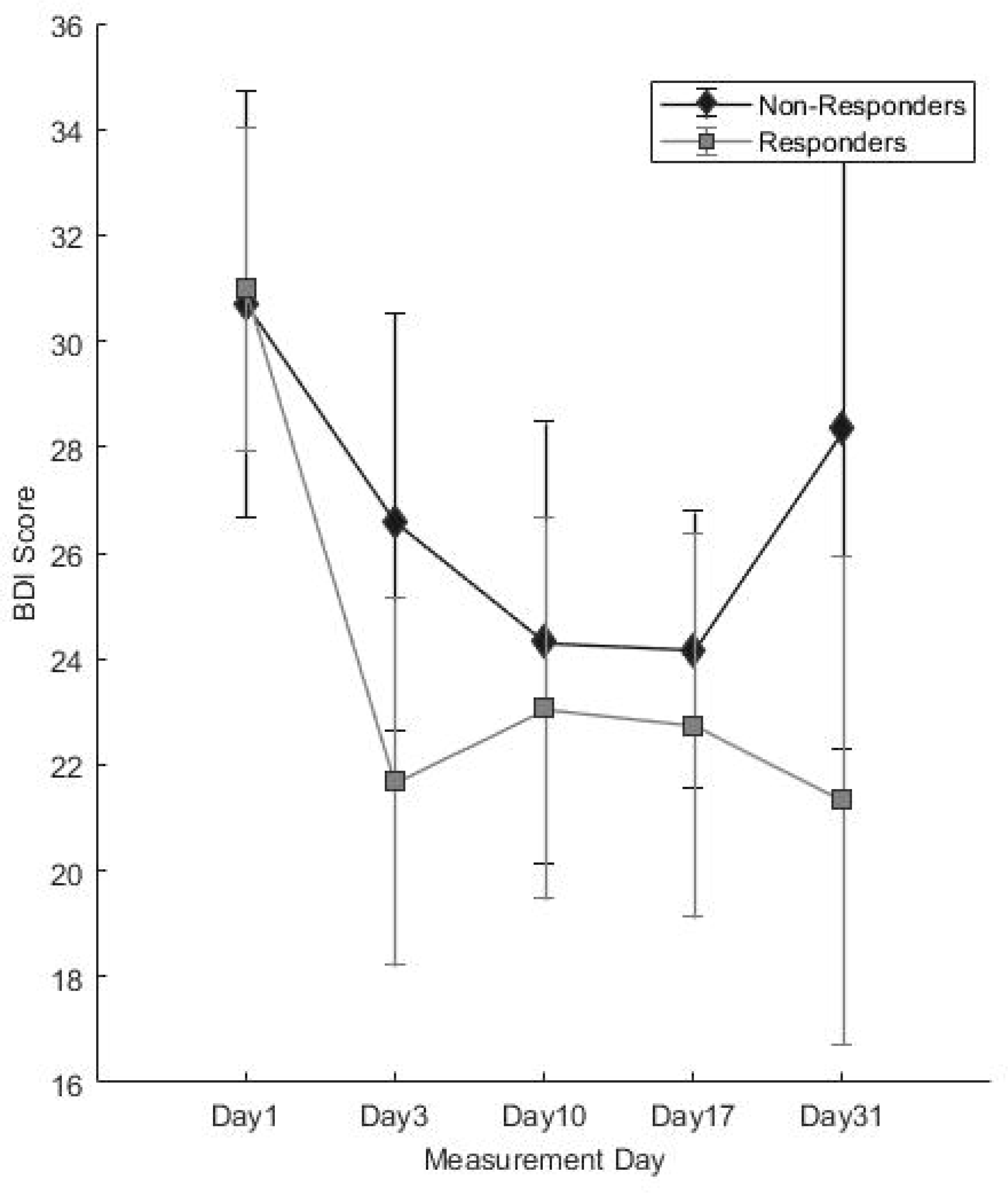
Post-treatment trajectories of a) MADRS and b) BDI-II scores. Error bars denote 95% confidence intervals. BDI-II = Beck Depression Inventory II; MADRS = Montgomery-Äsberg Depression Rating Scale.

For the *MADRS*, significant main effects were observed for *response* (*F*_1,68,573_ = 6.155, *p* = 0.016); *measurement day* (*F*_4,87,373_ = 49.388, *p* < 0.001); *measurement day*\**response* interaction (*F*_4,87.492_ = 5.339, *p* = 0.001); and *season* (*F*_3,61.090_ = 3.854, *p* = 0.014). *MADRS* scores showed a significant decrease on all measurement days compared to baseline (all *p* < 0.001) (Fig 4a, **Table S4b**). The interaction term revealed significantly lower scores in responders than non-responders on Days 4 (*t* = 4.242, *df* = 83.491, *p* < 0.001); 10 (*t* = 2.394, *df* = 80.704, *p* = 0.019); and 31(*t* = 2.767, *df* = 55.519, *p* = 0.008), but not Day 17 (*t* = 1.169, *df* = 81.646, *p* = 0.246). *MADRS* scores were significantly higher in spring than during other seasons (vs. summer *p* = 0.013; autumn *p* = 0.020; winter *p* = 0.002). No significant effects of *sex* (*p* = 0.420), *age* (*p* = 0.519), *AaO* (*p* = 0.855), *FH* (*p* = 0.784), or *diagnosis* (*p* = 0.850) or *PRS for MDD* (*p* = 0.155) were observed.

For *BDI-II*, significant main effects were observed for *measurement day* (*F*_4,65.719_ = 13.140, *p* < 0.001), *season* (*F*_3,57.224_ = 9.733, *p* < 0.001) and *sex* (*F*_1,56.431_ = 5.091, *p* = 0.028). *BDI-II* scores decreased significantly on all measurement days compared to baseline (all *p* < 0.001) (Fig 4b, **Table S4c**) and significantly higher in spring compared to all other seasons (all *p* < 0.001). No significant interaction between *response*\**measurement day* was observed (*F*_4,65.719_ = 65.719, *p* = 0.296, only a trend for higher scores in non-responders on Day 31, *p* = 0.085). Higher BDI-II scores were observed in women (*t* = 2.256, *df* = 56.431*p* = 0.28). No significant effects of *response* (*p* = 0.918), *age* (*p* = 0.960), *AaO* (*p* = 0.941), *FH* (*p* = 0.566), or *diagnosis* (*p* = 0.712) or *MDD PRS* (*p* = 0.559) were observed.

## Discussion

The observed association between response and both younger age at presentation (Clark *et al*, 2007; Pflug *et al*, 1971a) and higher age at disease onset (Rudolf *et al*, 1978) replicates previous reports. The finding that responders and non-responders did not differ in terms of baseline depressive symptom scores is consistent with reports of depression severity not influencing SD response (Clark *et al*, 2007; Pflug *et al*, 1971a; Szuba *et al*, 1991; van den Burg and van den Hoofdakker, 1975). Previously reported associations with diurnal variation were not observed (Haug, 1992; Martiny *et al*, 2013; Reinink *et al*, 1990).

In the present cohort the proportion of response to SD was on the higher end of the range reported in a recent meta-analysis, in which response rates ranged from 7-78% (Boland *et al*, 2017). The authors hypothesized that the small individual sample sizes were likely to contribute to this wide range of response rates. It is of note that the mean sample size of these studies was approximately 23 and approximately 66% of these studies had smaller sample sizes. In the present study, we applied the same protocol consistently in a large sample of patients over a protracted period of time, making the response rate we observed more robust and less prone to spurious factors which might be observed in small samples assessed during relatively short time spans.

We examined genetic burden for MDD using PRS, finding significantly higher scores in nonresponders than controls. We also found higher PRS in non-responders compared to responders, although differences were not statistically significant. These preliminary data suggest that underlying biological differences may be involved in SD effects and may suggest an avenue for exploration in larger samples.

Although initial depression severity did not differ in responders and non-responders, differing subjective mood and mood trajectories were observed. The main effect of better mood observed in responders may indicate better attitude towards the treatment, and should be further explored. Interestingly, both responders and non-responders experienced some degree of mood improvement during SD; although the interaction between response and timepoint was not statistically significant (Figure 3a, see also **Figure S1**), this might be qualitatively accounted for by mood scores responders in crossing the mid-point of the VAS (i.e. Fig 3a, from the ‘negative’ to ‘positive’ side of the scale). Further research should use multi-dimensional mood assessments to better examine the changes. Tiredness, previously reported to be a predictor of response (Bouhuys *et al*, 1995), did not differ between responders and non-responders except for in the early evening. On the whole, tiredness trajectories were similar in all participants.

Correlations observed between *BDI-II/MADRS* suggest validity of both scales. Although trajectories appeared similar, the interaction between response and assessment day was significant for *MADRS*, but not *BDI-II*. This may be attributable to 1) differences in number of items and points assigned to each item and 2) the fact that the *BDI-II* is a subjective measure, containing many items assessing maladaptive personality traits (Svanborg and Asberg, 2001) unlikely to change in the short-term. Interestingly, women reported higher *BDI-II* but not *MADRS* scores than men which may further suggest that the symptoms contributing to depression are different between the sexes.

Importantly, these longitudinal scores reflect clinical treatment outcomes, suggesting that response to SD may be a general indicator of response to further treatment. We included season to control for possible effects (daylight hours, temperature), finding more pronounced depressive symptoms in the spring, which is consistent with previous research showing exacerbation of mood disorders in spring (Cobb *et al*, 2014). We note that while the *BDI-II* and *MADRS* detected no baseline differences between groups, the VAS did. The VAS measures positive mood, which is not assessed in depressive symptom scales. This suggests that future studies should quantify positive mood, and as mentioned above, that measurement of the multiple dimensions of mood/affect would allow more rigorous characterization of behavioural patterns during SD.

This study had several limitations. First, as this was a naturalistic study, patients were not randomised/stratified with respect to medication, diagnoses, age at onset, or illness duration. Second, the sample size was too small to control for all potential influences, despite being one of the larger reported SD cohorts to date. Third, response to SD was assessed using the CGIC, which does not allow specification of which symptoms have changed. However, changes in both the *MADRS* and *BDI-II* scores were consistent with the CGIC (Figure 4). Fourth, comparison with a group of depressed patients not undergoing SD would have strengthened the interpretation of our findings. Finally, we did not correct *p*-values for multiple testing.

In conclusion, the rapid, pronounced effects of SD render it a well-controlled, efficient model (Dallaspezia and Benedetti, 2015). We propose that it is a promising context to apply novel methods such as genome-wide analyses (of the epi/genome and proteome) (Arnardottir *et al*, 2014; Bunney *et al*, 2015; Koike *et al*, 2012; Massart *et al*, 2014; Moller-Levet *et al*, 2013; Ramaker and Dulawa, 2017; Takahashi, 2017) and further more ecologically valid techniques such as ambulatory assessment (Trull and Ebner-Priemer, 2013). We believe that such an approach is suitable to not only link observed phenotypic changes with underlying biological factors, but to do so in a way such that depression heterogeneity (and interindividual differences) can be dissected.

## Contributors

MR, MD and SHW designed the study. NT, SCvH, MG and MD administered sleep deprivation and clinical assessments. NT and JCF analysed the data, interpreted results and with MR wrote the main manuscript. JF, SHW, FS, JT assisted with analyses, interpretation of results and writing of the manuscript. AFJ, UEP, MMN, MD and MR interpreted results, and edited and wrote the manuscript. All authors discussed findings and implications, commented on the manuscript, and approve of the final version. The MDD Working Group of the Psychiatric Genomics Consortium provided GWAS data and commented on the manuscript.

